# Domain Compatibility and Linker Design Dictate the Success of Chimeric Cellulase Engineering

**DOI:** 10.1101/2025.09.10.675174

**Authors:** Aditi Konar, Arijit Mondal, Sneha Sahu, Supratim Datta

## Abstract

Efficient conversion of lignocellulosic biomass into fermentable sugars remains a major challenge due to individual cellulases’ limited synergy and catalytic efficiency. Engineering chimeric enzymes provides a promising strategy to streamline biomass hydrolysis by combining complementary catalytic activities in a single protein, thereby enhancing efficiency and lowering process costs. In this study, we constructed chimeric cellulases by fusing a thermophilic GH1 β-glucosidase (*Ts*BG) with endoglucanases from the GH5 (*Bs*EG2) or GH9 (*Bl*EG) families through flexible peptide linkers. Constructs containing BsEG2 exhibited a pronounced loss of β-glucosidase activity and reduced endoglucanase activity, whereas substitution with the full-length BlEG restored dual functionality under identical design conditions. The optimized chimera (*Bl*EG+(G_4_S)_2_+*Ts*BG) demonstrated enhanced catalytic performance, with a 4.8-fold lower *K*_m_, a 1.7-fold higher *V*_max_, and an increased *k*_cat_ (from 1088 to 1454 s^-1^). The chimera also exhibited enhanced stability, retaining ∼10 % higher activity under elevated cellobiose (up to 300 mM) and >90 % specific activity in 2.5 M NaCl. Molecular dynamics simulations further revealed that activity loss in non-optimized constructs arose from C-terminal structural instability and steric clashes, underscoring the critical role of domain orientation and linker flexibility in chimera design. These findings establish a chimeric cellulase that integrates endoglucanase and β-glucosidase activities in a single polypeptide, offering a robust and cost-effective biocatalyst for lignocellulosic biomass conversion.

## Introduction

The rapid growth of the global population and dwindling fossil fuel reserves have intensified the demand for sustainable energy alternatives. Renewable biofuels, particularly those derived from lignocellulosic biomass, present a viable solution for decarbonizing transportation fuels. Lignocellulosic biomass, sourced from agricultural waste, energy crops, forestry, and marine environments, is abundant, renewable, and avoids competition with food supplies [1]. The conversion of lignocellulosic biomass into fermentable sugars relies on cellulases-enzymes that hydrolyze cellulose into glucose through a synergistic action [2]. Endoglucanases randomly cleave internal β-1,4-glycosidic bonds in cellulose chains, exoglucanases processively liberate cellobiose units from chain ends, and β-glucosidases further hydrolyze cellobiose to glucose [3]. This sequential action ensures efficient substrate degradation while mitigating product inhibition [4, 5]. However, the high cost of cellulase production remains a major bottleneck, driving efforts to engineer more efficient enzyme systems or reduce dosage requirements.

To address this challenge, chimeric cellulases – fusion proteins combining functional domains – have emerged as a promising strategy to enhance synergy and minimize enzyme loads [6]. Previous work has focused on fusing endoglucanase–exoglucanase, endoglucanase-xylanase, or exoglucanase–β-glucosidase pairs. But fewer studies have explored fusing processive endoglucanases (which directly produce cellobiose) with β-glucosidases. Such designs could streamline hydrolysis, reduce purification costs, and improve biochemical properties. This study explored a less-studied combination where a processive endoglucanase, which directly produces cellobiose from insoluble biomass, was fused to a β-glucosidase. Such chimeric enzymes could improve catalytic efficiency, reduce purification costs, and enhance biochemical properties.

While carbohydrate-binding modules (CBMs) are frequently fused to catalytic domains, synergistic cellulase chimeras remain underexplored [7, 8]. Notable examples include thermostable endo-exoglucanase fusion of *Ct*GH5-F194A and *Ct*GH1 from *Clostridium thermocellum* and chimeras of β-1,4-glucosidase and endoglucanase from *Paenibacillus* sp. [9, 10]. An endoglucanase-xylanase chimera has been reported, where endoglucanase Endo5A was fused with a xylanase Xyl11D to broaden the substrate range for hydrolysis [11]. Computational approaches such as SCHEMA have further advanced cellulase engineering by enabling *de novo* design of chimeric enzymes through sequence recombination [12]. For instance, Banerjee et al. used SCHEMA to generate a cellulase-like protein property from GH48 cellulase sequences, while Heinzelman et al. recombined fungal exoglucanases to enhance thermostability [13, 14].

Designing a functional chimeric protein requires careful optimization of three key parameters. Firstly, the selected enzymes should function optimally under similar reaction conditions at a common pH_opt_. Secondly, each enzyme’s N- or C-terminal positioning must be empirically determined. Thirdly, selecting rigid, flexible, or tailored linkers for maximal activity [15, 16]. Here, we present a systematic approach to engineer a novel chimera by fusing *Bacillus* sp. GH5 endoglucanase (*Bs*EG2) with *Thermococcus* sp. β-glucosidase (*Ts*BG) [17], followed by iterative optimization of domain arrangement and linker design. We also incorporated a two-domain GH9 endoglucanase, *Bl*EG from *Bacillus licheniformis*, combining its catalytic domain and CBM to enhance substrate affinity [18]. Supported by computational modelling and molecular dynamics simulations, our work demonstrates how strategic domain fusion can enhance cellulolytic efficiency, offering a scalable path toward cost-effective biofuel production.

## Materials and methods

### Chemicals

All chemicals utilized in this study were of reagent grade. DNA polymerase, Restriction endonucleases, restriction exonucleases, Antarctic phosphatase, and DNA ligase were purchased from NEB (Beverly, USA). Eurofins Genomics synthesized the primers (Hyderabad, India). DNA ladder, dNTP mix, and protein ladder were from Thermo Scientific (Mumbai, India). All chemicals and substrates, including *p*NPGlc, were sourced from Sigma Aldrich (St Louis, USA) and Merck (India). Plasmid miniprep and gel purification kits were procured from Qiagen (Hilden, Germany). Plasmid concentrations were measured using a TECAN Infinite^®^ 200 Pro (TECAN Trading AG, Switzerland). A 30 kDa cut-off AmiconUltra-50 membrane (Merck Millipore, Bangalore, India) was used for protein concentration.

### Cloning and Assembly

Gene-specific primers (Table S1) were designed using IDT OligoAnalyzer (IA, USA) to amplify the individual and chimeric genes. *In silico* cloning was verified using SnapGene (CA, USA). The open reading frame of endoglucanase *Bl*EG and endoglucanase *Bs*EG2 was extracted from *Bacillus licheniformis* and *Bacillus* sp., respectively. The strains were a kind gift from Prof. S. Venkata Mohan, BEES Lab, IICT Hyderabad. The synthetic gene corresponding to the β-glucosidase (*Ts*BG) from *Thermococcus* sp. was synthesized (Gene accession number BankIt1899421 BG Thermo KU867869) and assembled by Gene Art (Thermo Fisher Scientific, Waltham, USA) [17]. The cDNA template was PCR amplified by Q5 high-fidelity DNA polymerase. Genes were amplified with gene-specific forward and reverse primers and assembled via Gibson assembly with primers containing linker sequences. Each tagged with a 6x-His tag, the assembled genes were cloned into the pET21b (+) vector.

### Protein Expression and Purification

A 5 mL overnight culture of *E. coli* BL21(DE3) transformed with chimeric constructs in the pET21b vector was grown in Luria Bertani (LB) media supplemented with ampicillin (100 *μ*g/mL at 37 °C. This overnight culture was used to inoculate a 1 L secondary culture at a 1:100 dilution. When the culture reached the OD_600_ of 0.6, protein expression was induced with IPTG (Isopropyl β-D-1-thiogalactopyranoside), followed by incubation at 25 °C for 12 hours. Cells were harvested by centrifugation at 8000 × g for 10 minutes at 4 °C.

All purification steps were performed using an ÄKTA pure 25 M (Cytiva, Marlborough, MA) with UNICORN software version 6.3. The pellet was resuspended in lysis buffer containing 10 mM potassium phosphate buffer (pH 7.4), 40 mM imidazole, 500 mM NaCl, 1 mM PMSF (Phenylmethylsulfonyl fluoride), and 1.2 mg/mL lysozyme. Cells kept on ice were sonicated at 40 % amplitude for 10 cycles, each consisting of a 20-second pulse followed by a 1-minute rest. The lysate was centrifuged at 13000 × g for 40 minutes at 4 °C, and the supernatant was collected.

Protein purification was performed using a 5 mL His Trap™ HP column (GE Healthcare Life Sciences, Pittsburgh, USA) equilibrated with buffer A (10 mM potassium phosphate buffer (pH 7.4), 40 mM imidazole, 500 mM NaCl). Protein was eluted using a linear gradient from 0 to 100 % of buffer B (10 mM potassium phosphate buffer (pH 7.4) containing 500 mM imidazole, and 500 mM NaCl). Eluted protein was concentrated and desalted using a HiTrap™ desalting column (GE Healthcare Life Sciences, Pittsburgh, USA in 20 mM phosphate buffer (pH 7.3). Purity was assessed via 10 % SDS-PAGE, and protein concentration was determined by measuring absorbance at 280 nm, using extinction coefficients calculated via the modified Edelhoch and Gill/Von Hippel method available on the ExPASy ProtParam tool (Swiss Institute of Bioinformatics)[19].

### Co-expression with chaperones

To improve the solubility and yield of the chimeric proteins, co-expression was performed with different chaperone sets using plasmids pGro7, pKJE7, pTf16, pG-Tf2, and pG-KJE8 (DSS Takara, New Delhi, India). To evaluate the impact of chaperone co-expression, total protein content from each culture was quantified using a Bradford Assay [20]. Equal amounts (30 µg) of protein were analyzed by 10 % SDS PAGE, followed by western blotting using primary and secondary antibodies. Protein bands were visualized using the SuperSignal West Femto chemiluminescent substrate kit (Thermo Scientific, Mumbai, India), following the manufacturer’s protocol. The effect of co-expression on BL21(DE3) growth was monitored over 10 hours post-induction. Growth curves were normalized by the following equation:

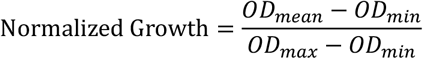

The values were scaled to between 0 and 1 for each time point.

### Determination of optimal pH and temperature

The optimal pH (pH_opt_) was determined by measuring the specific activity of endoglucanase on sodium carboxymethyl cellulose (CMC-Na, sourced from wood pulp, viscosity 1500-3000 cP, Na: 6.5-9.5 %) using the 3,5-dinitrosalicylic acid (DNS) assay and β-glucosidase on 4-nitrophenyl β-D-glucopyranoside (*p*NPGlc) across a pH range of 4.0–8.0 in McIlvaine buffer (0.2 M disodium hydrogen phosphate and 0.1 M citric acid). The temperature dependence of enzyme activity was assessed over a range of 35 °C to 80 °C, following incubation for 30 minutes (endoglucanase) and 5 minutes (β-glucosidase).

### Enzyme activity assays

All enzymatic reactions were performed in a thermomixer (Eppendorf, Hamburg, Germany). Endoglucanase activity was measured using sodium carboxymethyl cellulose as substrate with 0.05 μg of enzyme in McIlvaine buffer at respective pH_opt_ and T_opt_ in a final reaction volume of 150 μL for 20 minutes. The 3,5-dinitrosalicylic acid (DNS) assay was utilized to quantify reducing sugars, with glucose as the calibration standard. The β-glucosidase was assayed using the chromogenic substrate, *p*NPGlc, with 0.2 μg of enzyme in a 100 μL reaction in McIlvaine buffer at its pH_opt_ and T_opt_ for 5 minutes. The assay was stopped by adding 100 μL of glycine-NaOH buffer (pH 10.8). One unit of enzymatic activity is defined as the amount of enzyme required to release 1μ*μ*mol of product per minute from the substrate. All assays were performed in triplicate, and the standard deviation was reported.

Michaelis-Menten modelling using GraphPad Prism determined steady state kinetic parameters with values obtained by extrapolating the fitted curve. For the chimeric enzyme *Bl*EG+(G_4_S)_2_+*Ts*BG, the reaction rate was quantified by DNS assay, measuring total reducing sugars produced in 30 minutes with increasing CMC substrate concentrations. Due to the high viscosity of sodium carboxymethyl cellulose (CMC), substrate saturation could not be achieved [21, 22]. Hence, Michaelis–Menten kinetics were measured up to 26 mg/mL, and *K*_m_ and *V*_max_ were extrapolated from curve fitting.

### Half-life assays

The half-life of the chimeric enzymes was determined by incubating them at their optimal temperature and pH. Residual activity was measured at regular intervals by removing aliquots, centrifuging, and assaying specific activity. Data were fitted to a one-phase decay model in GraphPad Prism. The substrate used was CMC-Na for endoglucanase and *p*NPGlc for β-glucosidase. For each sample, blanks without enzymes were subtracted for any background absorbance.

### Cellobiose tolerance

The effect of cellobiose (0 to 400 mM) on endoglucanase activity was assessed using AzoCMC (Neogen, Cochin, India) as the substrate, under the optimum assay conditions. The specific activity of the endoglucanase part of the chimeric enzyme without any added cellobiose was considered 100 % and the relative specific activity was calculated.

### Salt tolerance

Salt tolerance was evaluated by measuring enzymatic activity in the presence of increasing NaCl concentrations. Reactions were conducted with CMC-Na and *p*NPGlc as a substrate for endoglucanase and β-glucosidase, respectively, at their pH_opt_ and T_opt_. The specific activity of salt-free chimeric enzymes was set as 100 %.

### Molecular Dynamics (MD) Simulation

MD simulations of the 3D structures of AlphaFold-modelled chimeric proteins were performed using GROMACS 2021.1. The modelled chimera was placed inside a cubic box, maintaining a minimum distance of 10 Å between any atom and the box edges. The CHARMM36m force field was employed, and the system was solvated using the CHARMM-modified TIP3P water model [23, 24]. Counterions were added to neutralize the system’s net charge. Energy minimization was performed using the steepest descent method for 50,000 steps in each case. This was followed by equilibration in two phases: first, a 100 ps equilibration under the NVT ensemble at 300 K; second, a 1 ns equilibration under the NPT ensemble at 1 bar to stabilize pressure. A 200 ns production run was then performed with a 2 fs integration time step. Simulation trajectories were analyzed using built-in GROMACS analysis modules.

### Synergy

Synergy reactions were performed by the parental proteins (*Bl*EG and *Ts*BG) added in a one-pot reaction in their free forms and were compared to the reaction by the chimeric protein *Bl*EG+ (G_4_S)_2_+ *Ts*BG. For the chimeric protein reaction, 1 μg of *Bl*EG+ (G_4_S)_2_+ *Ts*BG was added to the reaction mixture. For the cocktail reaction, a mixture of 0.6 μg BlEG and 0.4 μg *Ts*BG was used, reflecting the 60:40 molecular weight ratio of the chimeric protein. Reactions were carried out in McIlvaine buffer at the optimal pH and temperature for 90 minutes. Substrate options included 1% soluble sodium carboxymethyl cellulose (CMC) and 4 % insoluble Avicel. Reducing sugars released during the reaction were quantified using the DNS assay.

## Results and Discussion

### Selection of proteins for chimeric construct

The selection of proteins for constructing a chimeric enzyme is critical to ensure its functionality and efficiency. To engineer a novel chimeric enzyme, we fused the catalytic domain of a thermostable endoglucanase from glycoside hydrolase (GH) family 5 (GH5), *Bs*EG2 (manuscript under preparation), from *Bacillus* sp., with *Ts*BG, a thermophilic β-glucosidase from *Thermococcus* sp., a member of the GH1 family [17]. To link these domains, we employed a flexible (G_4_S)_2_ linker composed of glycine for flexibility and serine to prevent unwanted domain interactions via hydrogen bonding with water [25]. We also used a rigid linker, (EA_3_K)_2_, for comparison. We tested both N-terminal and C-terminal fusions of *Bs*EG2 with *Ts*BG to evaluate optimal orientation. Additionally, we substituted *Bs*EG2 with an alternative endoglucanase from *Bacillus licheniformis, Bl*EG, which possesses a GH9 catalytic domain and a CBM [18]. One variant was *Bl*EG at the N-terminus and *Ts*BG at the C-terminal, joined by a flexible linker (G_4_S)_2_, while the other variant contained a rigid linker (EA_3_K)_2_ to minimize steric hindrance between the catalytic domains [26]. All constructed chimeric cellulase variants are summarized in Figure 1.

**Figure 1.**
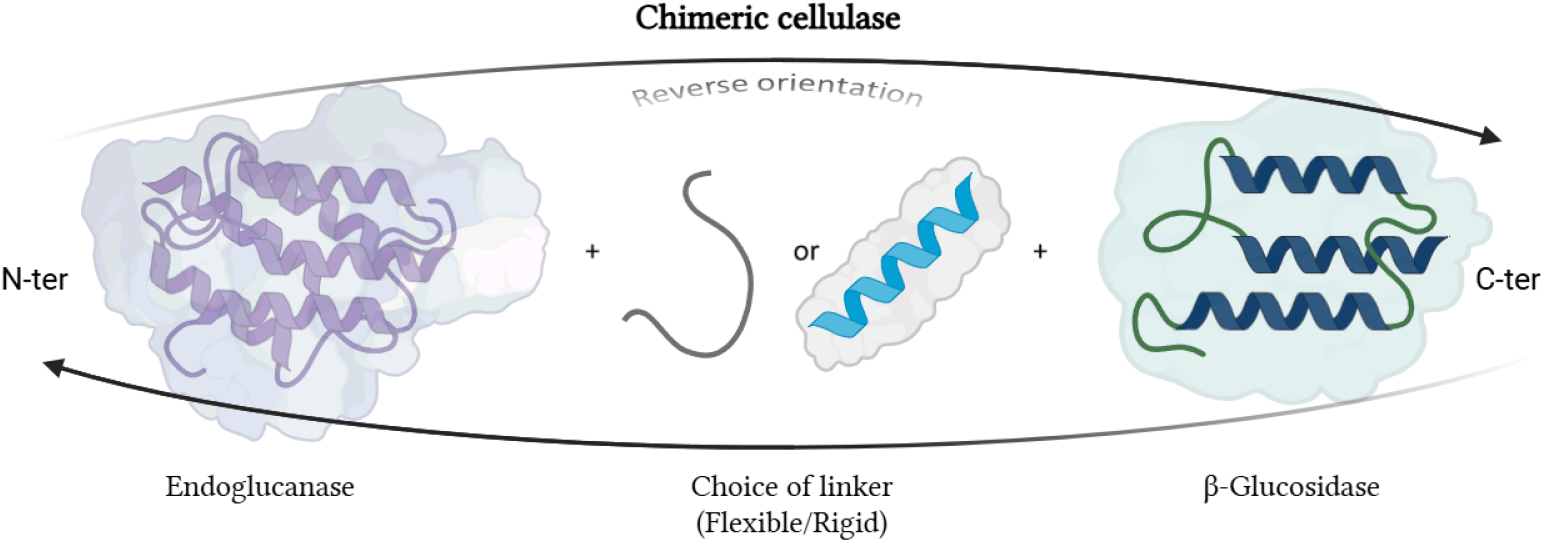
Schematic of the workflow for designing the optimal chimeric constructs made up of an endoglucanase, a linker (flexible and rigid), and a β-glucosidase

### Cloning, expression, and characterisation of chimeric proteins

The genes encoding the chimeric proteins were cloned into the pET-21b(+) vector under the control of a T7 promoter, with each construct containing a C-terminal 6×His tag for purification. The recombinant plasmids were then transformed into *Escherichia coli* BL21(DE3) cells for heterologous expression. Protein production was induced with isopropyl-β-D-thiogalactopyranoside (IPTG), and inducer optimization revealed 0.4 mM IPTG as optimal for maximal chimeric protein yield. Following induction, cells were harvested, and the His-tagged chimeric proteins were purified by Ni-NTA affinity chromatography. However, the yields of the chimeric proteins were lower than those of their parental forms. We employed chaperone-assisted co-expression to improve solubility and folding of the most promising construct, BlEG+(G4S)2 +TsBG. A panel of chaperone-encoding plasmids, including pGro7, pG-KJE8, pTf16, pGTf2, and pKJE7, was obtained (DSS Takara, New Delhi, India) and co-transformed individually into *E. coli* BL21(DE3) along with the chimeric construct. Among the tested systems, *Bl*EG+(G_4_S)_2_+*Ts*BG co-expression with the pGro7 plasmid, encoding GroEL and GroES, yielded the highest soluble protein levels, as confirmed by Western blot analysis using an anti-His antibody compared to the chimeric protein expressed without any chaperone (Figure S1). Notably, this co-expression strategy did not impair cell growth (Figure S2), indicating that the improved protein folding outweighed any additional metabolic burden from chaperone expression.

### Activity of the chimeric protein

The enzymatic assays were performed using 1 % CMC as the substrate for endoglucanase activity, and *p*NPGlc as the substrate for β-glucosidase activity. The chimeric enzymes exhibited optimal temperature (Figure 2) and pH (Figure S3) profiles identical to their parental enzymes (endoglucanase and β-glucosidase). Fusion of *Bs*EG2 to *Ts*BG via a (G_4_S)_2_ linker resulted in an 86 % reduction in β-glucosidase specific activity and a 15 % decrease in endoglucanase activity compared to the free parental enzyme. In contrast, the *Bl*EG-*Ts*BG chimera, constructed with the same flexible (G_4_S)_2_ linker, retained both endoglucanase and β-glucosidase activities, likely due to *Bl*EG’s CBM acting as a spacer between the catalytic domains of *Bl*EG and *Ts*BG. Among all the tested chimeras, *Bl*EG+(G_4_S)_2_+*Ts*BG displayed the highest activity, matching the activity of the parental enzymes. Consequently, this chimera was selected for further study.

**Figure 2.**
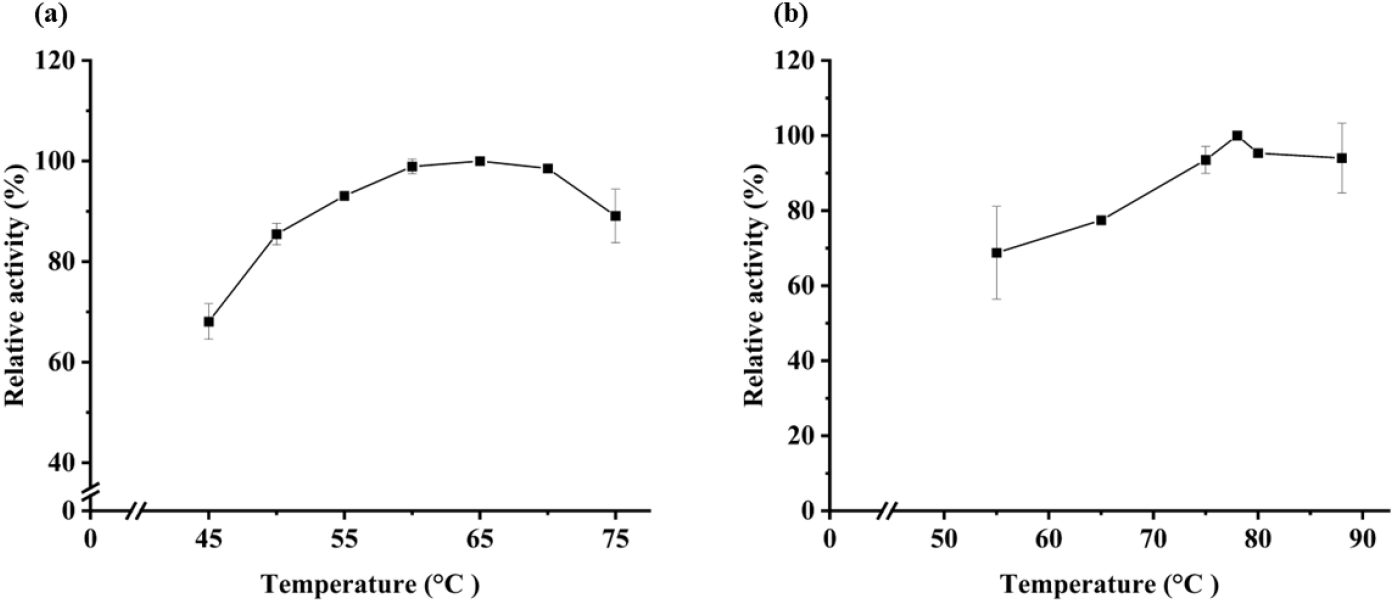
The temperature optima of *Bl*EG (a) and *Ts*BG (b) were determined under standard assay conditions. The optimal temperature of *Bl*EG (a) was determined by measuring activity on 1 % (w/v) carboxymethyl cellulose (CMC). The optimal temperature of *Ts*BG (b) was determined by measuring specific activity on 20 mM *p*-nitrophenyl-β-D-glucopyranoside (*p*NPGlc). Relative activity (%) is expressed as a percentage of the maximum observed activity (set to 100 %). Data represent mean values ± standard deviation (n ≥ 3 independent experiments performed in triplicate).

### Kinetic parameters of the chimeric protein

The kinetic parameters of free *Bl*EG (Uniport ID: H1AD14) have been previously reported [18]. For the chimera *Bl*EG+(G_4_S)_2_+*Ts*BG, we determined apparent *K*_*m*_, *V*_*max*_, and *k*_*cat*_ from endoglucanase activity assays (Figure 3). The *K*_*m*_ decreased from 32.4 mg/mL in the free parental enzyme to 6.7 mg/mL in the chimera, indicating enhanced substrate binding affinity. This improvement may arise from the fused β-glucosidase rapidly hydrolyzing cellobiose, reducing product inhibition and freeing the endoglucanase active site-a known bottleneck in cellulases like those from *Trichoderma reesei* [27]. Additionally, *V*_max_ increased from 306.2 to 524.4 µM min^-1^, and *k*_cat_ increased from 1088 to 1454 s^-1^, suggesting higher catalytic efficiency. These results highlight a potential advantage of chimeric enzymes over free enzyme cocktails, as the chimera exhibits both higher substrate turnover and stronger substrate affinity. However, chimeric protein design carries inherent variability in folding, which can impact activity and stability. For instance, studies on cellulase chimeras such as the combination of *Te*Egl5A and *Po*Cel5 have reported both improvements and a decrease in activity and stability [28]. In a library of ten GH5 cellulase chimeras made from a mesophilic (*Po*Cel5, T_opt_: 60 °C) and a thermophilic parent (*Te*Egl5A, T_opt_: 90 °C), only five were active. Some chimeras (H4-H6) showed reduced activity (17-51% compared to the parental enzyme) and stability, whereas others (H8, H9) exhibited markedly improved properties: temperature optima increased to 70–80 °C, T_50_ rose by 15–19 °C, T_m_ increased by 16.5–22.9 °C.

**Figure 3.**
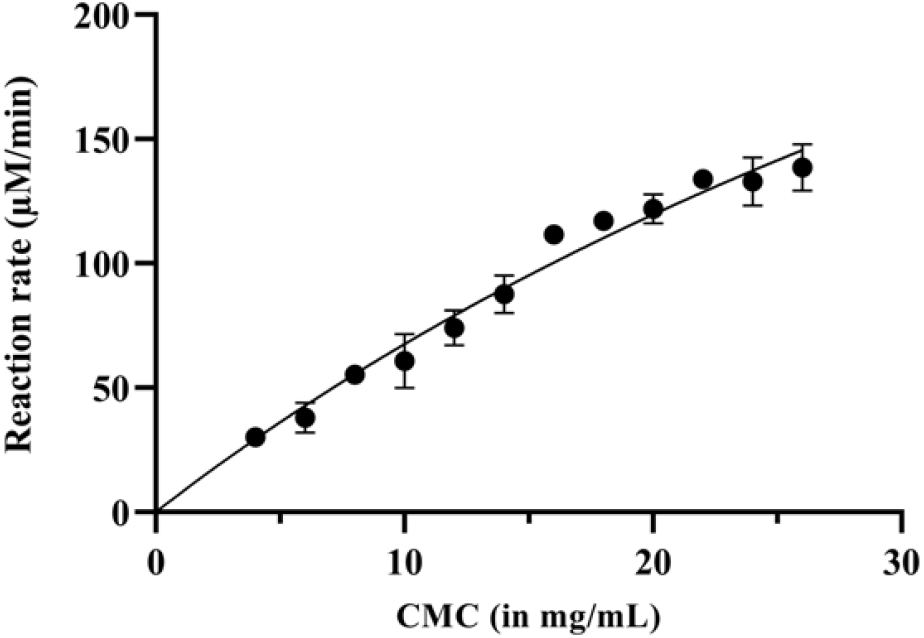
The Michaelis-Menten kinetics for *Bl*EG+(G_4_S)_2_+*Ts*BG was plotted, and the kinetic parameters estimated from the Michaelis-Menten assay were as follows: *K*_m_ = 6.7 mg/mL, *V*_max_ = 524.4 µM min^-1^, and *k*_cat_ =1454 s^-1^, respectively. Errors represent the standard deviations of independent reactions.

### Thermal stability of the chimeric proteins

The half-lives of *Bs*EG2 are 10 days at 55 °C, *Ts*BG is 55 minutes at 78 °C, and BlEG is 14 days at 65 °C [17, 18]. In *Bs*EG2+ (G_4_S)_2_+ *Ts*BG, the half-life of *Ts*BG increased up to 115 minutes (Figure S4). However, the half-life of the endoglucanase in both *Bs*EG2+(G_4_S)_2_+ *Ts*BG and *Bl*EG+(G_4_S)_2_+ *Ts*BG decreased significantly. Specifically, the half-life of *Bl*EG at 65 °C dropped to 2.6 hours, while *Ts*BG’s half-life at 78 °C remained near 60 minutes (Figure 4). This reduction in stability is consistent with previous reports on GH9 endoglucanase (Cel9A) chimeric constructs featuring different CBMs [7]. The observed decrease in half-life highlights a common challenge in engineering multi-domain chimeric proteins: the alteration of folding dynamics and inter-domain instability.

**Figure 4.**
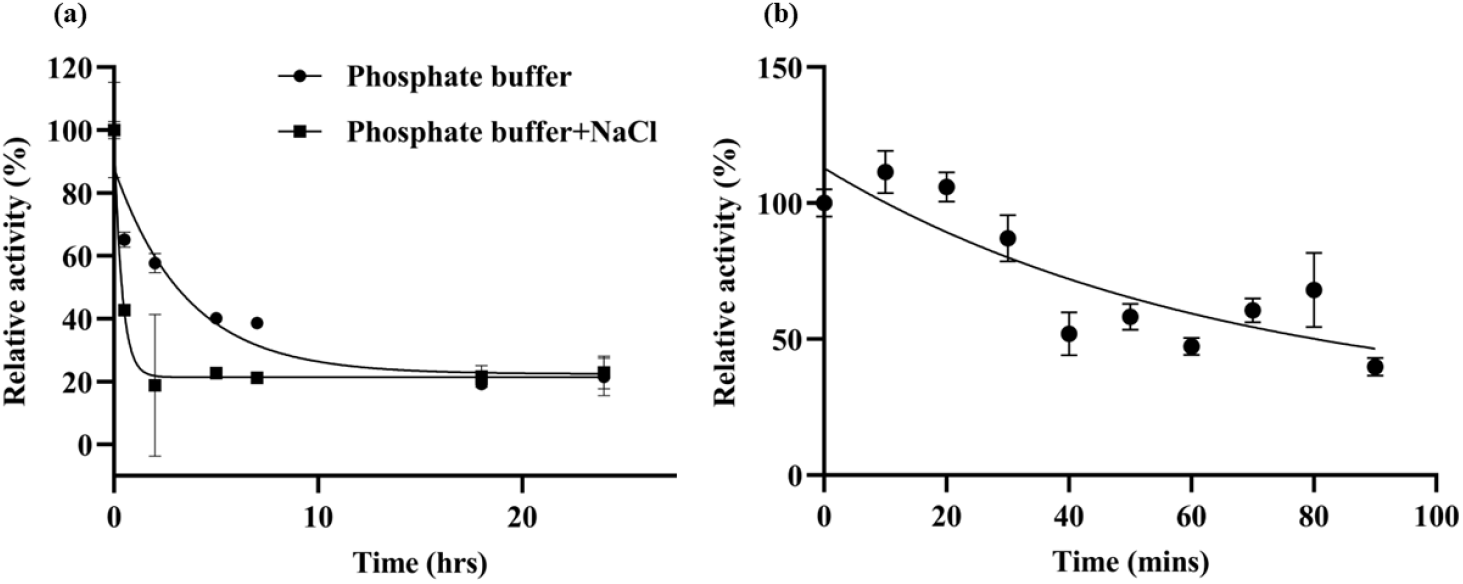
Half-life of chimeric proteins: (a) The half-life of *Bl*EG was measured by determining enzyme activity at its optimal temperature of 65 °C for up to 25 hours. The protein was incubated in 20 mM phosphate buffer, pH 6.5, and 20 mM phosphate buffer with 75 mM NaCl, pH 6.5. (b) The half-life of *Ts*BG was measured in phosphate buffer, pH 6.5, at its optimal temperature of 78 °C for up to 90 minutes. Half-life values were determined using a one-phase decay fit in GraphPad Prism. Error bars represent the standard deviation of independent experiments.

### Effect of cellobiose feedback inhibition on endoglucanase activity

Cellobiose, the primary product of *Bl*EG, inhibits endoglucanases [29]. In the presence of increasing cellobiose concentrations from 0 to 300 mM, a drop in specific activity of *Bl*EG in the *Bl*EG +(G_4_S)_2_ +*Ts*BG chimera was observed to be almost 10 % less than that of the free parental protein at each cellobiose concentration (Figure 5) suggesting that the fused β-glucosidase efficiently hydrolyzes cellobiose, reducing its accumulation and mitigating feedback inhibition of the endoglucanase. In the chimera, the proximity of endoglucanase and β-glucosidase enables immediate utilization of cellobiose released from the endoglucanase active site by the β-glucosidase, offering a key advantage over free enzyme cocktails where diffusion delays hinder such synergy.

**Figure 5.**
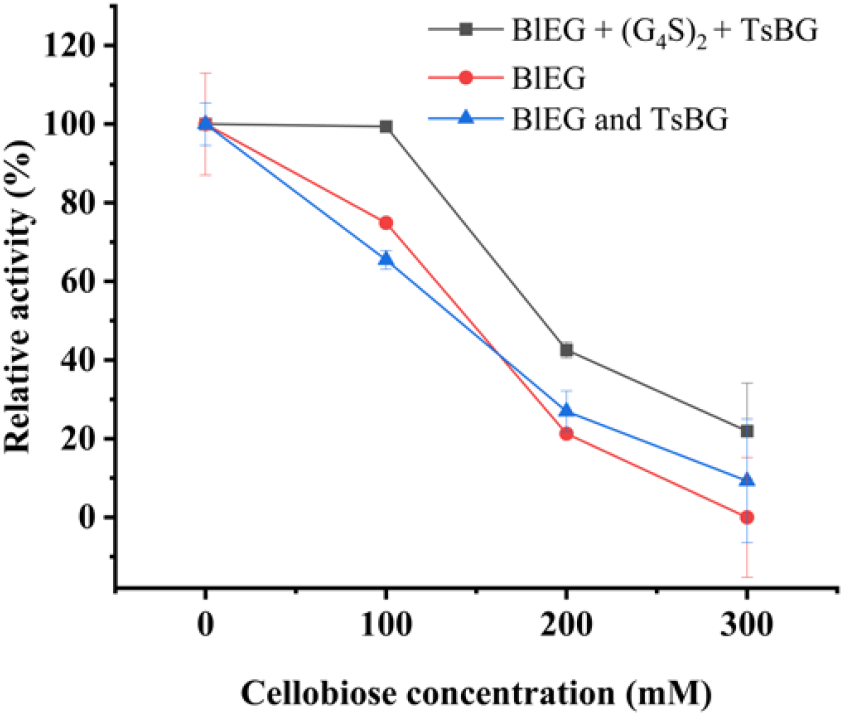
The cellobiose tolerance of *Bl*EG in the chimeric protein *Bl*EG+(G_4_S)_2_+*Ts*BG was compared to its free parental form, *Bl*EG, and a cocktail of *Bl*EG and *Ts*BG across various cellobiose concentrations, up to 300 mM. Error bars represent the standard deviation of independent experiments.

### Effect of salt (NaCl) on the chimeric protein

Using seawater and marine biomasses as sustainable resources is promising, but high salt concentrations in marine-derived cellulose and seawater often impair enzyme efficiency. To assess NaCl tolerance, we measured the relative activity of *Bl*EG and *Ts*BG in the chimera, *Bl*EG + (G_4_S)_2_+ *Ts*BG. Remarkably, *Bl*EG in *Bl*EG + (G_4_S)_2_+ *Ts*BG retained over 90-95 % specific activity at 2.5 M NaCl, whereas the free parental enzyme *Bl*EG (UniProt ID: H1AD14) retained only 75-80 % specific activity [18]. This halotolerance likely stems from *Bl*EG’s net negative surface charge conferred by acidic amino acid residues that stabilize the protein under high salinity [30]. Intriguingly, *Ts*BG’s relative specific activity increased with salt concentration (Figure 6), possibly due to salt-induced stabilization of misfolded and native protein conformations, preventing the aggregation of the active site pockets [31]. Consistent with prior findings, *Ts*BG (Uniport ID: O08324) not only maintained activity at elevated salt levels but also exhibited an extended half-life [17], aligning with published reports of other chimeric cellulases [32].

**Figure 6.**
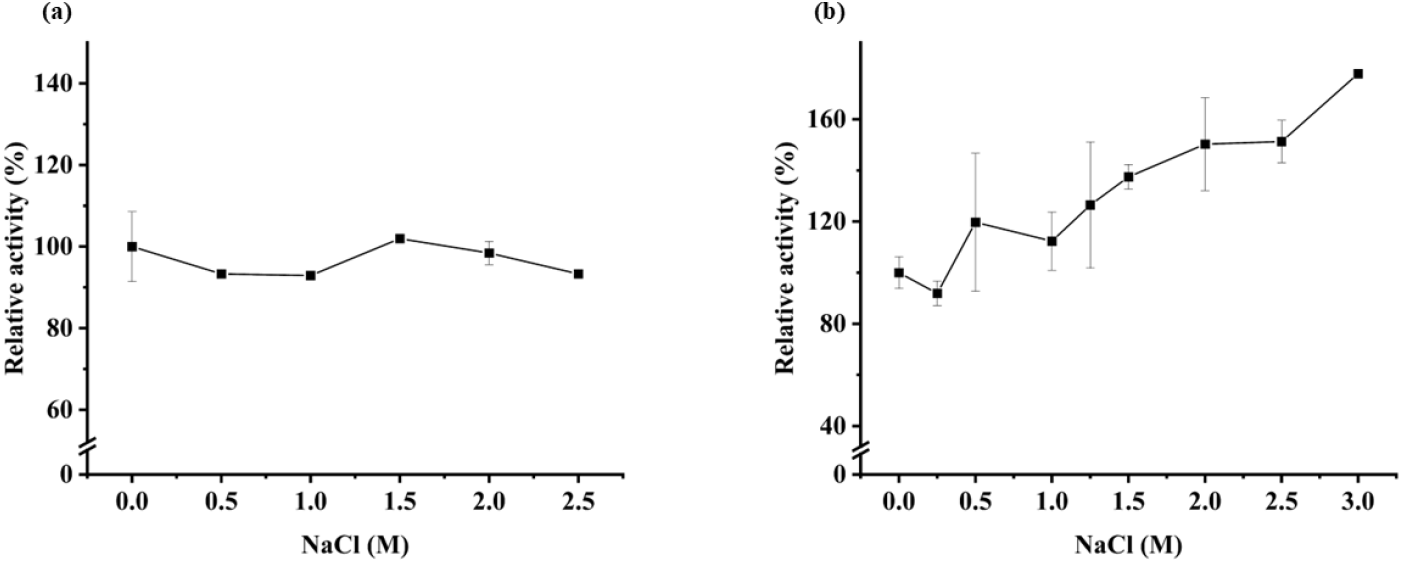
The NaCl tolerance of *Bl*EG (a) and *Ts*BG (b) in the chimeric protein *Bl*EG+(G_4_S)_2_+*Ts*BG was evaluated across varying NaCl concentrations, up to 2.5 M. Error bars represent the standard deviation of independent experiments.

### Structural determinants of chimera functionality

MD simulations were conducted on all four designed chimeras to further investigate the structural basis for the functional differences among the chimeric constructs, particularly focusing on the dynamics of the two enzyme components. We assessed structural deviations and flexibility over the 200 ns trajectories using root mean square deviation (RMSD; Figure S5) and root mean square fluctuation (RMSF; Figure S6) analyses. Notably, the functional chimeras *Bs*EG2+(G_4_S)_2_+*Ts*BG and *Bl*EG+(G_4_S)_2_+*Ts*BG exhibited stable dynamics, with both the N- and C-terminal domains maintaining RMSD values within approximately 5 Å. In contrast, the non-functional constructs *Ts*BG+(G_4_S)_2_+*Bs*EG2 and *Bl*EG+(EA_3_K)_2_+*Ts*BG displayed stable N-terminal domains (RMSD within ∼5 Å) but significantly higher C-terminal RMSD values (ranging from 7.5 to 20 Å), indicating pronounced structural instability.

To further explore the interdomain interactions in the functional chimeras, we employed *gmx mindist*, which calculates the minimum distance (the shortest atomic distance between any pair of atoms from the two catalytic domains) and the number of contacts (defined as atomic pairs lying within 7 Å)[33]. Analysis revealed that among the functional constructs, *Bl*EG+(G_4_S)_2_+*Ts*BG, which retained full β-glucosidase and endoglucanase activity, maintained a 3-to 4-fold greater minimum interdomain distance compared to *Bs*EG2+(G_4_S)_2_+*Ts*BG, which exhibited over 85 % loss of β-glucosidase activity. The increased interdomain distance resulted in fewer interatomic contacts and reduced steric clashes (Figure 7). In contrast, the approximately 3 Å proximity in *Bs*EG2+(G_4_S)_2_+*Ts*BG likely caused steric hindrance, restricting conformational flexibility and impairing activity.

**Figure 7.**
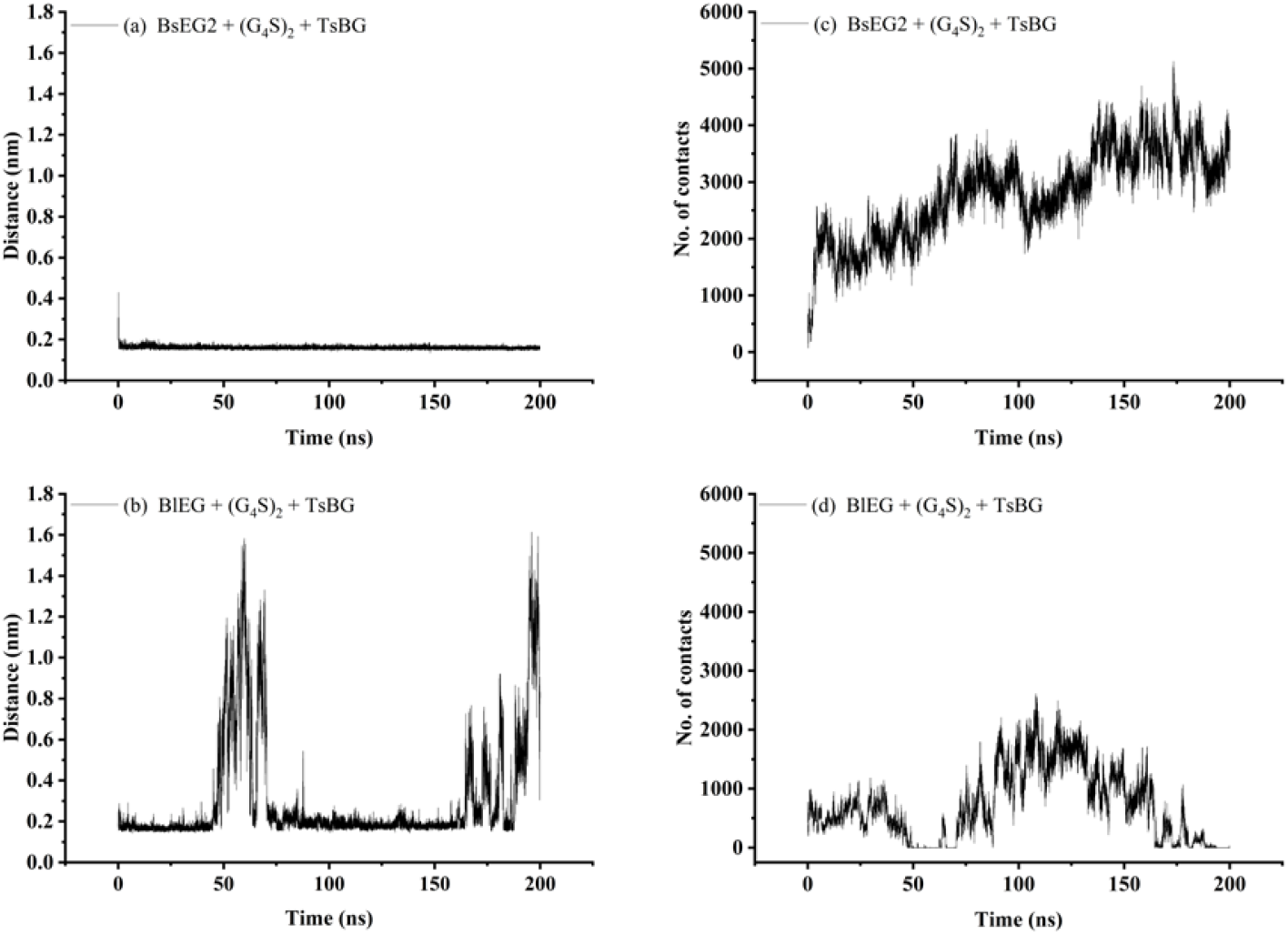
Interdomain minimum distance and contact analysis of functional chimeric constructs (from MD). Minimum interdomain distance profiles, defined as the shortest atomic distance between the two catalytic domains, are shown for *Bs*EG2+(G_4_S)_2_+*Ts*BG (a) and *Bl*EG+(G_4_S)_2_+*Ts*BG (b). Interdomain contact profiles, defined as the number of atomic pairs within 0.7 nm, are shown for *Bs*EG2+(G_4_S)_2_+*Ts*BG (c) and *Bl*EG+(G_4_S)_2_+TsBG (d).

Overall, the MD trajectories highlighted two distinct features. The non-functional constructs exhibited pronounced C-terminal instability, where excessive flexibility, characterised by high RMSD, has been previously reported to promote unfolded state populations, biasing the overall fold toward partially unfolded conformations [22, 34]. In the functional constructs, interdomain spacing was crucial in determining whether dual activity was preserved. These computational insights could be integrated into chimera design pipelines to more efficiently predict domain compatibility and guide experimental efforts.

### Synergistic Activity

In this study, we utilized a processive endoglucanase that primarily facilitates the conversion of long cellulosic chains into cellobiose units. Following this, β-glucosidase acts to hydrolyze cellobiose into glucose. We measured enzyme synergy by assessing activity on CMC (soluble substrate) and Avicel (insoluble substrate). The combination of *Bl*EG+(G_4_S)_2_+*Ts*BG produced 85-100 % reducing sugars, achieving levels comparable to those produced by the individual enzymes in a synergistic cocktail. This demonstrates its potential to replicate native enzyme synergy within a single fusion construct. (Figure 8).

**Figure 8.**
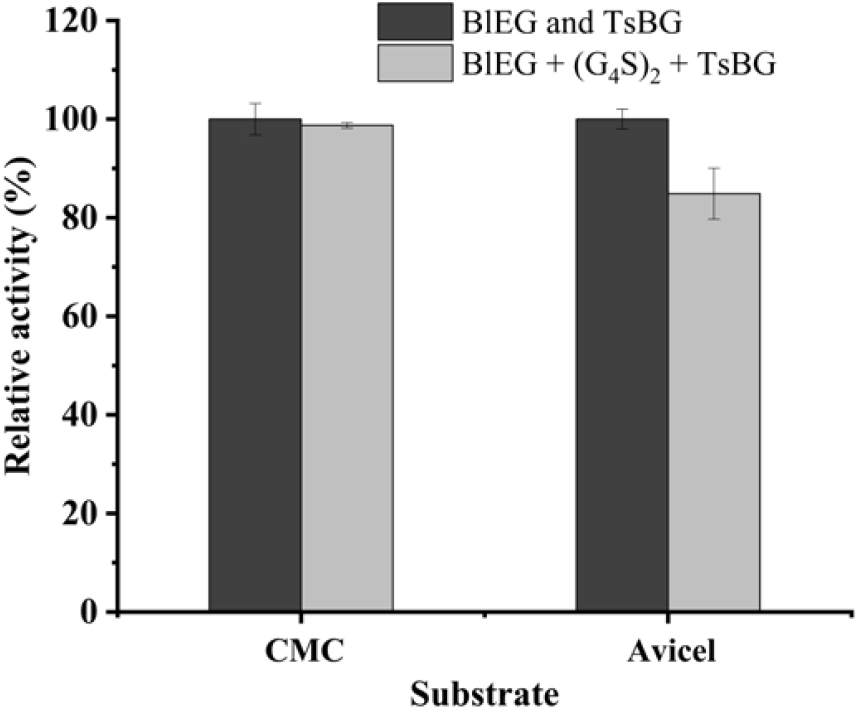
The synergistic action of *Bl*EG and *Ts*BG was analyzed both in their free enzyme forms and as a chimeric fusion, *Bl*EG + (G_4_S)_2_ + *Ts*BG, on both soluble substrate (1 % CMC) and insoluble substrate (4% Avicel). Reaction was carried out for 90 minutes at their T_opt_ and pH_opt_, and the reducing sugars generated were quantified by DNS assay. Error bars represent the standard deviation of independent experiments.

## Conclusions

The chimera *Bl*EG+(G_4_S)_2_+*Ts*BG emerged as the top performer, with the CBM in *Bl*EG optimally positioning the catalytic domain of *Ts*BG. The flexible (G_4_S)_2_ linker minimizes folding interference, enhancing overall functionality. MD simulations confirmed its structural stability and optimal interdomain spacing, mitigating steric clashes and enabling efficient substrate processing. Biochemical characterization revealed that *Bl*EG+(G_4_S)_2_+*Ts*BG exhibited enhanced halotolerance and product stability, along with superior kinetic performance. Specifically, there was a 79 % reduction in *K*_m_, a 71% increase in *V*_max_, and a 34 % increase in *k*_cat_ compared to the free *Bl*EG. These improvements indicate a substantial boost in catalytic efficiency and product yields, demonstrating a synergistic one-pot reaction similar to that of separately added enzymes. This engineered chimera represents an efficient, salt-tolerant biocatalyst for biomass degradation, showcasing significant potential for industrial applications that require robust enzymatic performance.

## Acknowledgements

This work was supported by the Science & Engineering Research Board (SERB), Government of India, CRG/2023/002111 (SD) and Department of Biotechnology (DBT), Government of India, BT/PR47801/BCE/8/1812/2023, and IISER Kolkata Academic Research Funds to CCES, IISER Kolkata. AK is supported by a Senior Research Fellowship from CSIR, Government of India, and SS is supported by a IPh.D. fellowship from IISER Kolkata. A KVPY fellowship by the Department of Science and Technology, Government of India, supported AM. We sincerely thank Prof. Gautam Basu (IISER Kolkata) for his invaluable suggestions.

## Tables

**Table 1.**
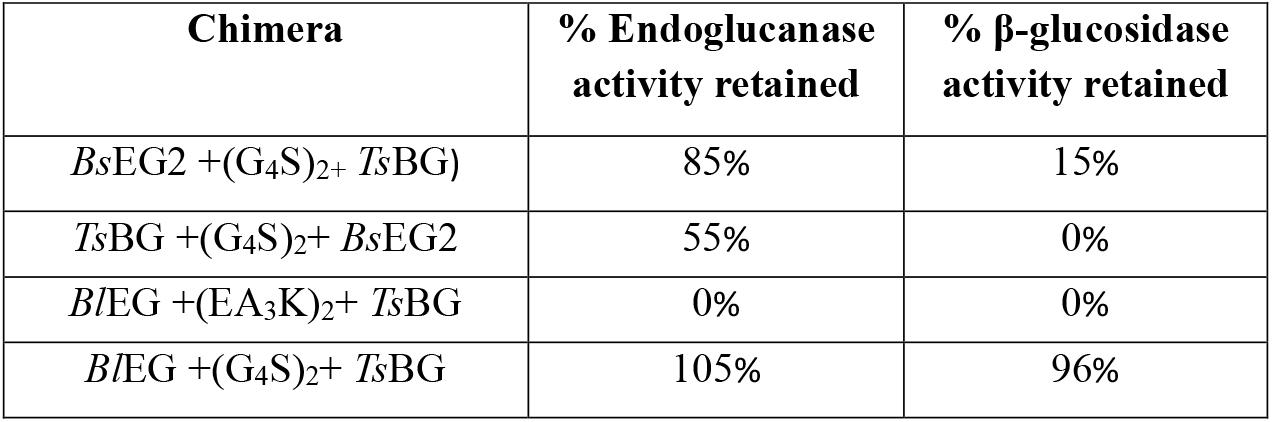
The % activity retained by the chimeric enzymes *Bs*EG2 +(G_4_S)_2_+ *Ts*BG, *Ts*BG +(G_4_S)_2_+ *Bs*EG2, *Bl*EG +(EA_3_K)_2_+ *Ts*BG, *Bl*EG +(G_4_S)_2_+ *Ts*BG in their chimeric forms, as compared to their respective activities in free forms, *Bl*EG (H1AD14), *Ts*BG (O08324), and *Bs*EG2 [17, 18].

