## Supplementary Files for "Domain Compatibility and Linker Design Dictate the Success of Chimeric Cellulase Engineering"

**Table S1.** Sequences of the primers used for cloning

| **Primer names** | **Primer sequences** |
|---|---|
| *Bl*EG EcoRI fwd primer | ACGGAATTCATGGCGGACATTGTATTTCGGCAAGGC |
| *Bl*EG $(G_4S)_2$ rev primer | GCCACCGCTGCCACCGCCACCgtcgggctcatttccaaaaacaagccg |
| *Ts*BG $(G_4S)_2$ fwd primer | GGCAGCGGTGGCGGTGGCAGCATGTTTCGTTTTCCGGATGGATTCC |
| *Ts*BG XhoI rev primer | CCGCGGCTAGTGGTGATGATGGTGGTGGTGGTGGTGGTGCTCG |

**Figure S1**. Western blot analysis of chimera, *Bl*EG+$(G_4S)_2$+*Ts*BG, expression. Lanes: L1, pGro7; L2, PG-KJE8; L3, uninduced; L4, IPTG induction only; L5, PTf16; L6, PGTf2; L7, pKJE7; L8, SDS page ruler. All samples were induced with 1 mM IPTG. Additional inducers used: arabinose (0.5 mg/mL) for L1, L2, L5, and L7; tetracycline (5 ng/mL) for L1 and L6.

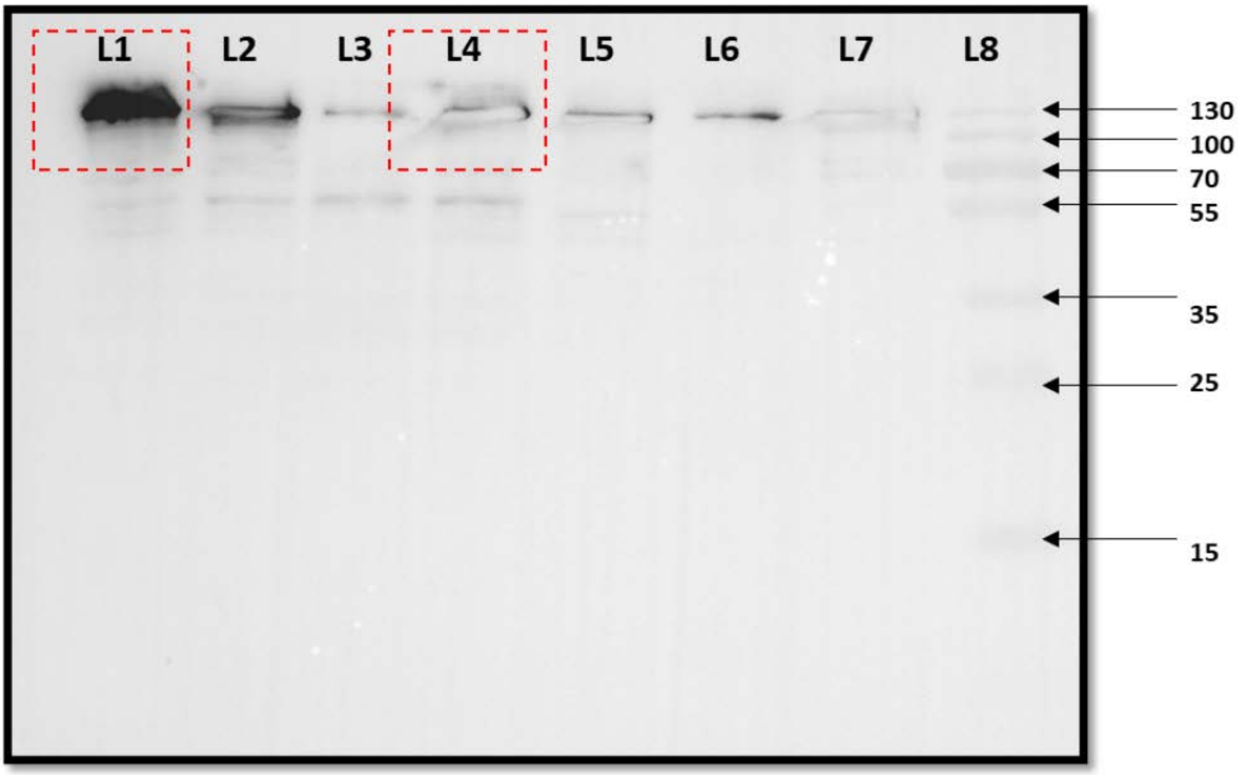

**Figure S2**. Growth curve analysis of *E. coli* cells expressing the chimeric protein *Bl*EG+$(G_4S)_2$+*Ts*BG (black) compared to co-expressing with the chaperone combination GroEL and GroES from plasmid pGro7 (red). Cells were cultured for 10 hours at 37°C, and $OD_{600}$ measurements were taken at regular intervals.

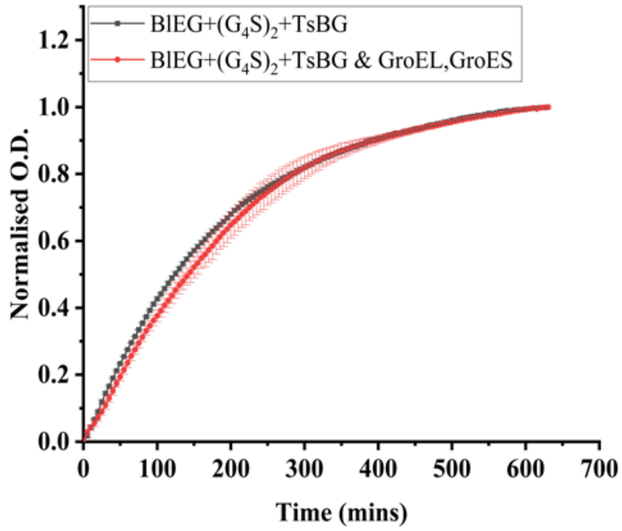

**Figure S3**. The effect of pH was determined by incubating the chimeric protein *Bl*EG+$(G_4S)_2$+*Ts*BG in McIlvaine buffer between the pH range of 4–8, at its optimal temperature for 30 minutes to measure the % relative activity.

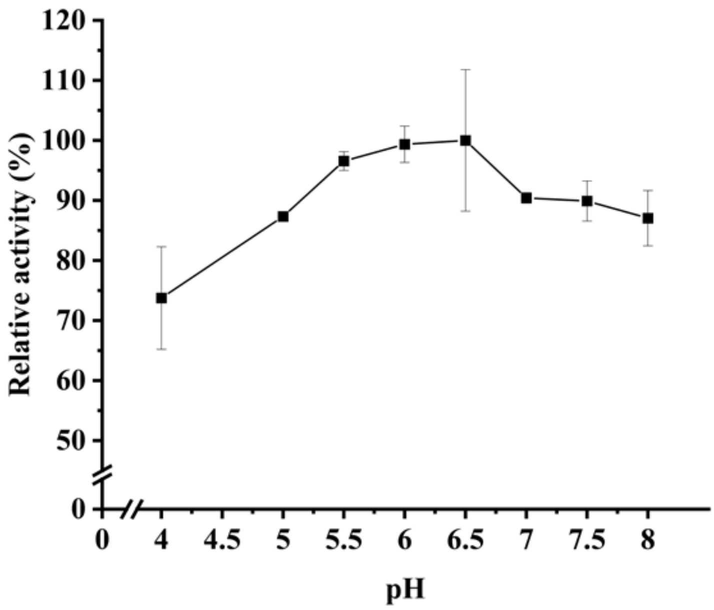

**Figure S4**. Half-life determination of *Ts*BG in the chimeric form *Bs*EG2+($G_4S$)$_2$+*Ts*BG. The half-life of *Ts*BG was measured at its optimal temperature (78 °C) up to 120 minutes, and the % relative activity was calculated by setting the initial activity (at 0 minutes) to 100 %.

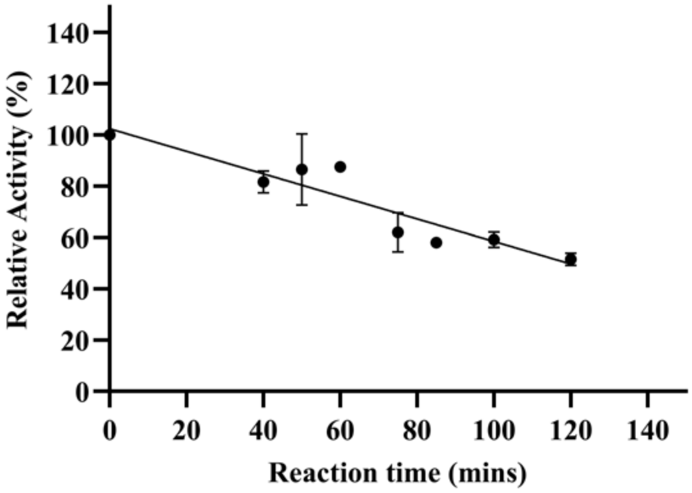

**Figure S5**. Root mean square deviation (RMSD) profiles of chimeric constructs, *Bs*EG2+$(G_4S)_2$+*Ts*BG (a), *Ts*BG+$(G_4S)_2$+*Bs*EG2 (b), *Bl*EG+$(G_4S)_2$+*Ts*BG (c), and *Bl*EG+$(EA_3K)_2$+*Ts*BG (d). Backbone RMSD values over MD trajectories for all four chimeras, showing overall structural stability and domain-specific deviations. The plot tracks the root-mean-square deviation (RMSD) of atomic positions relative to the starting structure, with separate plots for different regions of each chimeric protein: the N-terminal domain (black) such as *Bs*EG2 (a), *Ts*BG (b), *Bl*EG (c,d); the linker (either $(G_4S)_2$ or $(EA_3K)_2$) connecting the domains (red), and the C-terminal domain (blue) such as *Ts*BG (a), *Bs*EG2 (b), *Ts*BG (c,d).

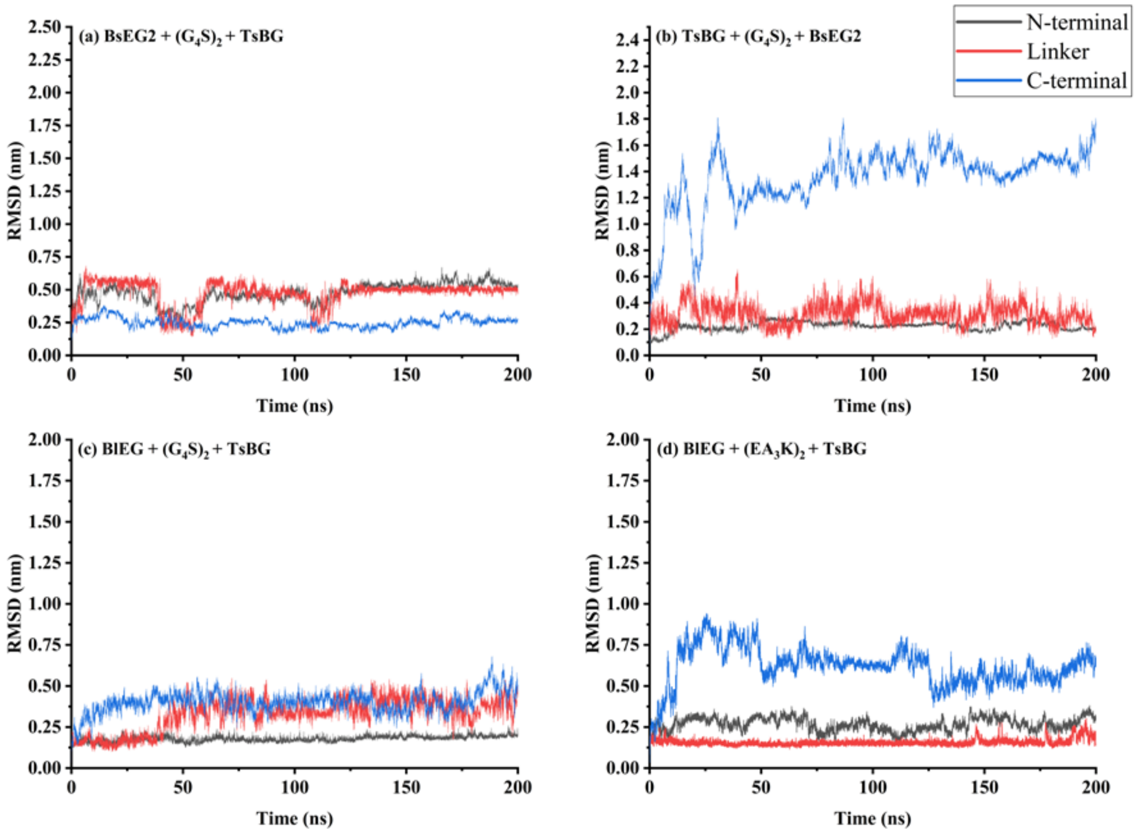

**Figure S6.** Root mean square fluctuation (RMSF) profiles of chimeric constructs. $C_\alpha$ RMSF values over 200 ns MD trajectories for each of the four chimeraic proteins, highlighting domain-wise flexibility. The plot shows residue-level fluctuations around their average positions during the simulation, with separate curves for different regions of each chimeric protein: the N-terminal domain (black) such as *Bs*EG2 (a), *Ts*BG (b), *Bl*EG (c,d); the linker (either $(G_4S)_2$ or $(EA_3K)_2$) connecting the domains (red), and the C-terminal domain (blue) such as *Ts*BG (a), *Bs*EG2 (b), *Ts*BG (c,d).

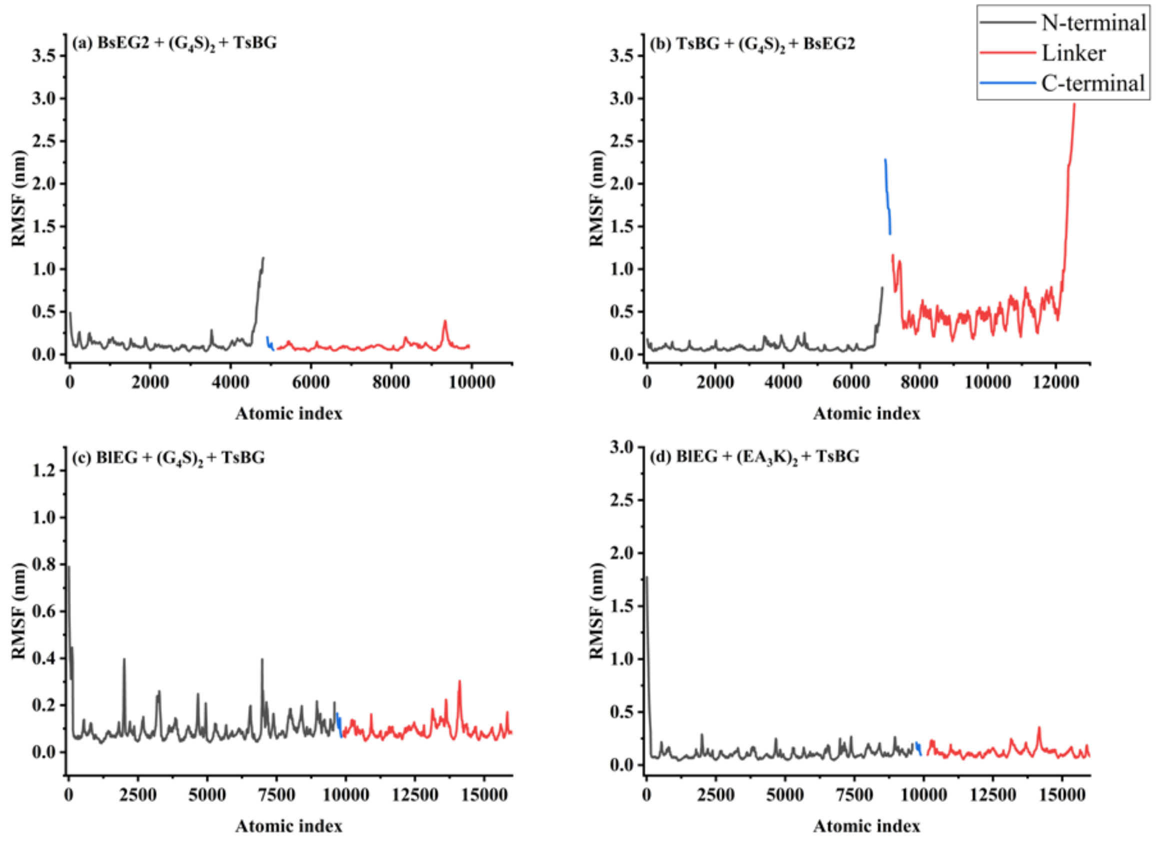
